# Heterogeneous nuclear ribonucleoprotein K is overexpressed in acute myeloid leukemia and causes myeloproliferative disease in mice via altered *Runx1* splicing

**DOI:** 10.1101/2021.02.05.429385

**Authors:** Marisa J. L. Aitken, Prerna Malaney, Xiaorui Zhang, Shelley M. Herbrich, Lauren Chan, Huaxian Ma, Rodrigo Jacamo, Ruizhi Duan, Todd M. Link, Steven M. Kornblau, Rashmi Kanagal-Shamanna, Carlos E. Bueso-Ramos, Sean M. Post

**Author notes:** To whom correspondence should be addressed, **Corresponding Author:** Sean M. Post, Ph.D., Department of Leukemia, The University of Texas MD Anderson Cancer Center 1515 Holcombe, Blvd, Houston, TX 77030, USA. These authors contributed equally (Marisa J. L. Aitken, Prerna Malaney).

## Abstract

Acute myeloid leukemia (AML) is driven by numerous molecular events that contribute to disease progression. Herein, we identified hnRNP K overexpression as a recurrent abnormality in AML that is associated with inferior patient outcomes. In murine hematopoietic stem and progenitor cells, hnRNP K overexpression altered self-renewal and differentiation potential. Furthermore, murine transplantation models revealed that hnRNP K overexpression resulted in fatal myeloproliferative phenotypes. Using unbiased approaches, we discovered a direct relationship between hnRNP K and *RUNX1*—a master transcriptional regulator of hematopoiesis often dysregulated in leukemia. Molecular analyses revealed hnRNP K-dependent alternative splicing of *RUNX1*, resulting in the generation of a functionally distinct isoform. Taken together, we have established hnRNP K as an oncogene in myeloid leukemia that binds *RUNX1* RNA, altering its splicing and subsequent transcriptional activity. These findings shed new light on a mechanism of myeloid leukemogenesis, paving the way for new drug discovery efforts.

**Highlights:**

- hnRNP K, an RNA binding protein, is overexpressed in AML and correlates with poor clinical outcomes
- hnRNP K overexpression in murine HSPCs drives fatal myeloproliferative phenotypes in mice
- hnRNP K’s oncogenicity can be attributed, at least in part, to its ability to bind and influence the splicing of the *RUNX1* transcript

## Introduction

Acute myeloid leukemia (AML) is an often devastating hematologic malignancy wherein normal hematopoiesis is superseded by rapid proliferation of incompletely differentiated myeloid cells. Taken as a whole, younger patients (<60 years) with AML have a 5-year overall survival of approximately 40%, while older patients tend to have much worse outcomes, with 5-year OS verging on 10% (Dohner et al., 2015; Juliusson et al., 2009; Slovak et al., 2000). Identification of recurrent genomic events in AML (e.g.; mutations in *FLT3 or IDH1/2*) has led to development of targeted therapeutic agents that improve patient outcomes and quality of life; however, many patients lack these genomic alterations, rendering them ineligible for such treatments. Furthermore, despite prolonging life, patients treated with these agents are often still at risk for relapse (DiNardo et al., 2018; Perl et al., 2019; Stein et al., 2019; Stone et al., 2017), highlighting the need to understand the molecular underpinnings of AML such that alternative, effective therapeutic options can be developed.

Myeloid malignancies, including AML and its frequently associated precursor condition myelodysplastic syndrome (MDS), often have alterations in RNA-binding proteins (RBPs). Splicing factors such as SRSF2, SF3B1, and U2AF1 are widely known RBPs in this context, and are often mutated (Graubert et al., 2011; Papaemmanuil et al., 2011; Yoshida et al., 2011). Importantly, other RBPs such as MUSASHI2, METTL3 are also aberrantly expressed, though infrequently mutated, in hematologic malignancies, and have been identified as critical to the pathogenesis of AML (Kharas et al., 2010b; Saha et al., 2019; Vu et al., 2017; Wang et al., 2019). Drugs targeting splicing or other RNA-binding properties of these proteins have been developed with therapeutic intent (Minuesa et al., 2019; Seiler et al., 2018). However, the roles of other RBPs in myeloid malignancies, including AML, have not been extensively deciphered.

An RBP of accumulating interest in hematologic and solid malignancies is heterogeneous nuclear ribonucleoprotein K (hnRNP K). Overexpression of hnRNP K has been associated with adverse pathology in a handful of small clinical studies evaluating solid tumors (Barboro et al., 2009; Carpenter et al., 2006; Chen et al., 2010; Wen et al., 2010; Zhou et al., 2010). Further, elevated hnRNP K expression is not uncommon in B-cell lymphoma and has been defined as an oncogene in this setting (Gallardo et al., 2019). These findings led to consideration that hnRNP K aberrancies may contribute to a broader array of hematologic malignancies. Because the expression and role of hnRNP K in AML is less clear, we sought to evaluate hnRNP K in this context.

Aberrations of RBPs often impact the expression of genes or proteins implicated in hematologic malignancies. For instance, RUNX1 is a crucial hematopoietic transcription factor with several isoforms that emerge due to alternative promoter usage and alternative splicing (Ghozi et al., 1996; Komeno et al., 2014; Miyoshi et al., 1995; Tanaka et al., 1995). *RUNX1* is a common target of translocations or mutations in leukemias, including AML (De Braekeleer et al., 2011; Osato, 2004; Osato et al., 1999). Interestingly, expression of different *RUNX1* isoforms arising via alternative splicing, specifically around exon 6, also alter hematopoiesis (Ghanem et al., 2018; Komeno et al., 2014; Sun et al., 2020).

In this study, we address the hypothesis that hnRNP K is overexpressed in AML, impacts clinic outcomes, and that this overexpression contributes to myeloid aberrations in a murine model. We find that hnRNP K overexpression leads to extramedullary hematopoiesis, gross hematopoietic abnormalities, and premature death in mice. Mechanistically, we identified that hnRNP K alters *RUNX1* splicing via its RNA-binding properties. hnRNP K interacts with *RUNX1* RNA in a sequence-specific manner, in humans and mice, and causes exclusion of *RUNX1* exon 6 (*RUNX1ΔEx6*), ultimately leading to expression of a more stable RUNX1 isoform with altered transcriptional activity. Further, expression of RUNX1ΔEx6 recapitulates an *in vitro* phenotype associated with hnRNP K overexpression, supporting the notion that hnRNP K mediates its hematopoietic alterations, at least in part, via altered *RUNX1* splicing. Finally, we identify that the RNA-binding KH3 domain of hnRNP K is a critical mediator of this *RUNX1* splicing event.

Taken together, these studies identify hnRNP K as a potential driver alteration in AML, nominating this protein as a putative drug target. In addition, hnRNP K-mediated altered splicing of *RUNX1* provides an alternate mechanism whereby RUNX1 is altered in AML in the absence of mutations or translocations. Finally, the methods used in this manuscript and the identification of specific domains of hnRNP K implicated in this splicing alteration provide a valuable set of tools to develop drugs that specifically disrupt hnRNP K-RNA interactions.

## Methods

### Reverse phase protein array (RPPA) data

Data is publicly available at www.leukemiaatlas.org (Hu et al., 2019). hnRNP K protein expression was compared to the median of healthy donor CD34+ bone marrow specimens.

### Plasmids

For generation of stable cell lines, lentiviral plasmids containing full-length hnRNP K, hnRNP K domain deletions, full-length RUNX1, or RUNX1ΔEx6 cDNA were cloned into the XhoI/NotI sites in the all-in-one tetracycline inducible lentiviral vector TRE3G-ORF-P2A-eGFP-PGK-Tet3G-bsd (TLO2026, transOMIC Technologies, Huntsville, AL). For retroviral plasmids used in fetal liver cells, the MSCV-AML1/ETO-IRES-GFP plasmid was obtained from Addgene (Addgene plasmid #60832; http://n2t.net/addgene:60832; RRID:Addgene_60832 (Zuber et al., 2009)) and *AML1/ETO* cDNA replaced with *HNRNPK, HNRNPKΔKH3, RUNX1(b)*, or *RUNX1ΔEx6* amplified from cDNA obtained from 293T cells and the pINDUCER-21-RUNX1 plasmid (Addgene plasmid #97043; http://n2t.net/addgene:97043; RRID:Addgene_97043 (Sugimura et al., 2017)), respectively. Cloning was done in XhoI/EcoRI sites in the MSCV-IRES-GFP plasmid. For generation of stably knocked down cell lines, tetracycline-inducible human HNRNPK PGK-TurboRFP shRNAs were purchased from Dharmacon (clone ID: V3IHSPGR_10844995 mature antisense TCGACGAGGGCTCATATCA, targeting exon 10), and two targeting the 3’ UTR, referred to as shHNRNPK ex16-1 (clone ID: V3IHSPGR_5114248, mature antisense AAGACACTAGAGCAAATTG) and shHNRNPK ex16-2 (clone ID: V3IHSPGR_9103684, mature antisense ATAAAATCCACTCACTCTG), and control PGK-TurboRFP (VSC11656, mature antisense TGGTTTACATGTTGTGTGA; Lafayette, CO).

For transient transfections, hnRNP K domain deletions (amplified with nested PCR (Ho et al., 1989)), *RUNX1(b)*, or *RUNX1ΔEx6* were ligated into XhoI/EcoRI restriction sites in the c-Flag pcDNA3vector (Addgene plasmid #20011; http://n2t.net/addgene:20011; RRID:Addgene_20011) (Sanjabi et al., 2005). Reporter assays to assess RUNX1 transcriptional activity were done using the pMCSF-R-luc plasmid (Addgene plasmid #12420; http://n2t.net/addgene:12420; RRID:Addgene_12420) (Zhang et al., 1994). The pCMV β-galactosidase plasmid was a gift from Dr. Vrushank Davé (University of South Florida).

### Stable cell line generation

Cell lines were spun for 90 minutes at 600 x g with filtered viral supernatant from 293T cells transfected with indicated plasmids (see above), and pCMV-VSV-G/pCMV-dR8.2 for human cell lines (Addgene plasmid #8454; http://n2t.net/addgene:8454; RRID:Addgene_8454 and Addgene plasmid #8455; http://n2t.net/addgene:8455; RRID:Addgene_8455, respectively (Stewart et al., 2003)) or pCL-ECO for fetal liver cells (Addgene plasmid #12371; http://n2t.net/addgene:12371; RRID:Addgene_12371 (Naviaux et al., 1996)). Human cells were selected in antibiotic (puromycin or blasticidin, Fisher Scientific, Waltham, MA) and appropriate fluorescence >90% required prior to downstream assays. Cells were maintained in strict tetracycline-free conditions until induction with 0.2 μg/mL doxycycline (Sigma-Aldrich, St. Louis, MO) for shRNA and 0.4 μg/mL doxycycline for overexpression. HSPCs were sorted for GFP positivity 72 hours after infection.

### HSPC isolation, transduction, and transplantation

All mouse studies were performed with approval from the Institutional Animal Care and Use Committee at MD Anderson under protocol 0000787-RN01/2. Pregnant wildtype CD45.2+ C57Bl/6 females were euthanized via CO_2_ exposure at day 13.5 of gestation, fetal livers dissected, and disrupted on a 70 μm filter into single-cell suspension. Cells were briefly subjected to red blood cell lysis (BDPharm Lyse, BD Biosciences, San Jose, CA) and resuspended in medium containing 37% DMEM (Corning Inc, Corning, NY), 37% Iscove’s modified Dulbecco’s Medium (Corning Inc, Corning, NY), 20% fetal bovine serum, 2% L-glutamine (200mM; Corning Inc, Corning, NY), 100 U/mL penicillin/streptomycin (Sigma-Aldrich, St. Louis, MO), 5×10^−5^ M 2-mercaptoethanol (Sigma-Aldrich, St. Louis, MO), recombinant murine IL-3 (0.2 ng/mL), IL-6 (2 ng/mL), and SCF (20 ng/mL; Stem Cell Technologies, Vancouver, BC) at high density overnight at 37°C prior to retroviral transduction (Schmitt et al., 2002; Zuber et al., 2009). For transplantation assays, NOD-scid-IL2R-gamma (NSG) mice were irradiated with 2.5 Gy prior to injection of 50,000 sorted cells into the retro-orbital sinus.

### Immunoblotting

Cells were homogenized in NP40 lysis buffer and standard immunoblotting procedures performed as previously described (Gallardo et al., 2019) using antibodies against hnRNP K (3C2), RUNX1 (EPR3099, both from Abcam, Cambridge, MA), β-actin (AC-15, Santa Cruz Biotechnology, Dallas, TX), Flag (F1804, Sigma Aldrich), HSP90 (ADI-SPA-836-D, Enzo Life Sciences)

### Colony formation assay

GFP-sorted HSPCs were cultured in Methocult (M3434, StemCell Technologies, Vancouver, BC). Colonies were counted after 7 days then gently disrupted in PBS and cells counted subjected to cytospin or flow cytometry.

### Flow cytometry

Cells were pre-treated with murine Fc block (TruStain FcX, BioLegend, San Diego, CA,) at room temperature for 15 minutes then incubated with antibodies Gr1 [RB6-8C5], CD11b [M1/70;], CD117 [2B8], CD45 [30F11] (all from BD Biosciences, East Rutherford, NJ,), and Sca-1 [D7; eBioscience, San Diego, CA,]. Flow cytometry was performed on a Gallios flow cytometer (Beckman Coulter, Brea, CA,). Data was analyzed using FlowJo (Beckton Dickinson, Franklin Lakes, NJ).

### Tissue harvest

Spleen, liver, and sternum were collected and immediately fixed in 10% neutral-buffered formalin. Paraffin-embedded blocks were sectioned and stained with standard hematoxylin/eosin. Cells were also collected from femurs and a portion of the spleen in a single cell suspension, subjected to RBC lysis, and processed for western blotting or qRT-PCR as described herein.

### Immunohistochemistry

Formalin-fixed paraffin-embedded tissues were deparaffinized in xylene and rehydrated in an alcohol gradient. Following antigen retrieval with citrate (pH 6.0), slides were incubated with 3% hydrogen peroxide/methanol prior to incubation with primary antibody at 4°C overnight. Primary antibodies: hnRNP K [3C2], CD3 [SP162], MPO [ab9535], CD14 [4B4F12] (Abcam, Cambridge, MA), CD34 [MEC14.7], CD117 [2B8] (ThermoFisher Scientific, Waltham, MA). Antibody-protein interactions were visualized with Vectastain Elite ABC and DAB peroxidase substrate kits (Vector Laboratories, Burlingame, CA) and counterstained with nuclear fast red.

### Peripheral blood analysis

Complete blood counts were performed with an ABX Pentra analyzer (Horiba, Kyoto, Japan). Peripheral blood smears were stained with Wright-Giemsa.

### RNA-Sequencing

RNA was extracted and purified from GFP+ sorted HSPCs using Zymo Quick-RNA columns (Zymo Research, Irvine, CA). Barcoded, Illumina-stranded total RNA libraries were prepared using the TruSeq Stranded Total RNA Sample Preparation Kit (Illumina, San Diego, CA). Briefly, 250ng of DNase I-treated total RNA was depleted of cytoplasmic and mitochondrial ribosomal RNA (rRNA) using Ribo-Zero Gold (Illumina, San Diego, CA). After purification, RNA was fragmented using divalent cations and double stranded cDNA was synthesized using random primers. The ends of the resulting double stranded cDNA fragments were repaired, 5′-phosphorylated, 3’-A tailed and Illumina-specific indexed adapters are were ligated. The products were purified and enriched by 12 cycles of PCR to create the final cDNA library. The libraries were quantified by qPCR and assessed for size distribution using the 4200 TapeStation High Sensitivity D1000 ScreenTape (Agilent Technologies, Santa Clara, CA) then multiplexed 3 libraries per lane and sequenced on the Illumina HiSeq4000 sequencer (Illumina, San Diego, CA) using the 75 bp paired end format.

### RNA-Seq analysis

Fastq files were pseudoaligned using Kallisto v0.44.0 (Bray et al., 2016) with 30 bootstrap samples to a transcriptome index based on the *Mus musculus* GRCm38 release (Ensembl). The resulting abundance data was further analyzed with Sleuth v0.30.0 (Pimentel et al., 2017) using models with covariates for both batch and condition. Gene-level 80 abundance estimates were calculated as the sum of transcripts per million (TPM) estimates of all transcripts mapped to a given gene. Wald tests were performed at a gene level for the “condition” covariate with a significance threshold of FDR <10%.

### fRIP analysis

Previously published data deposited to the Gene Expression Omnibus (GSE126479) was cross-referenced with known tumor suppressors and oncogenes.

### Identification of putative hnRNP K binding sites

As described previously (Gallardo et al., 2019), a computer-based algorithm was used that scan transcripts of interest for two or more (U/C)CCC motifs within 19 nucleotides.

### Fluorescence anisotropy (FA)

Recombinant hnRNP K protein, produced in E.coli as described previously (Gallardo et al., 2019), was serially diluted in PBS (0.1 nM to 10 μM) and incubated with 6-FAM labelled RNA oligonucleotides. FA values were measured with excitation wavelength 485 nm and emission wavelength 528 nm on a Synergy Neo multi-mode plate reader (BioTek, Winooski, VT). Data was fit to the following equation: FA=FAi+Bmax*[oligo]/(Kd+[oligo]) where initial FA is represented by FAi and the overall change in FA is represented by Bmax. Oligos: hRUNX1 5’UTR: CGCCCCCCCCCACCCCCCGCAGUAAUAAAGGCCCCUGA, hRUNX1 5’UTR(mut) CGCGCGCGCGCACGCGCCGCAGUAAUAAAGGCGCCUGA, hRUNX1 int5-6 UCUCUUCCCUCCCUCCUUCCCUCCCCCCAU, hRUNX1 int5-6(mut) UCUGUUCGCUCGCUCGUUCGCUCGCGCCAU, mRunx1 int5-6 UCCUCCUCCCUUCCCCUCCCGGUCCCUA, mRunx1 int5-6(mut) UCCUCCUCGCUUCGCCUCGCGGUCGCUA.

### Thermal shift assays

Recombinant hnRNP K protein (Gallardo et al., 2019) was incubated with SYPRO orange dye in the presence of DNA oligos [the DNA equivalent to RNA oligos used in FA assays]. Samples were heated from 25°C to 99°C in PBS buffer and fluorescence measured at each temperature increment using a StepOne Plus Real Time PCR System (Applied Biosystem, Foster City, CA). Negative controls with no protein were run on each plate. The first derivative of fluorescence was calculated at each temperature and the temperature corresponding to the minima was designated as the melting temperature of the sample.

### RT-PCR for RUNX1 isoforms

Equal amount of RNA per sample was converted to cDNA with iScript (BioRad, Hercules, CA). Standard PCR conditions were used to amplify cDNA using a BioRad Thermocycler at 95°C x 3 minutes, followed by 35 cycles of 95°C for 1 minute, 60°C for 1 minute, and 72°C for 3 minutes, and finally 72°C for 5 minutes. After amplification, equal amounts of PCR products were run on a 2% agarose gel with ethidium bromide and visualized using a Syngene G:Box EF2 gel doc system with GENESys image capture software. Primers (5’ to 3’): hRUNX1 (exon 5) forward GAAGTGGAAGAGGGAAAAGCTTCA, hRUNX1 (exon 7) reverse GCACGTCCAGGTGAAATGCG, hPPIA forward CCCACCGTGTTCTTCGACATT, hPPIA reverse GGACCCGTATGCTTTAGGATGA, mRunx1 (exon 5) forward CACTCTGACCATCACCGTCTT, mRunx1 (exon 7) reverse GGATCCCAGGTACTGGTAGGA, mPPIA forward GAGCTGTTTGCAGACAAAGTTC, mPPIA reverse CCCTGGCACATGAATCCTGG.

### Sanger Sequencing

DNA was purified from agarose gels using a gel purification kit (Invitrogen, Carlsbad, CA). Sequencing was performed on an ABI 3730XL sequencer using BigDye terminator cycle sequencing chemistry with the forward primer used in the RT-PCR reaction. For validation, another sequencing run was performed with the reverse primer used in the RT-PCR reaction. Data was provided as text files and chromatograms.

### Clinical RUNX1 isoform expression analysis

RNA-seq expression data for the full-length (ENST00000344691.8) and ΔEx6 (ENST00000399240.5) *RUNX1* isoforms and corresponding clinical data for 350 patients with AML from the BEAT AML 1.0 cohort was downloaded through the Vizome data portal (https://vizome.org/) February 2020 (Tyner et al., 2018).

### Protein stability assays

293T cells stably transduced with tetracycline-inducible constructs to overexpress *RUNX1*, either full-length or lacking exon 6. 400 ng/mL doxycycline was added to cells prior to the addition of cycloheximide (10 μM, Sigma-Aldrich, St. Louis, MO) with or without MG-132 (10 μM, SelleckChem, Houston, TX) for 1-8 hours. Cells were collected and lysed in NP40 lysis buffer with protease and phosphatase inhibitors prior to western blot.

### Transfections

293T cells were plated at identical confluencies and transfected using equal amounts of DNA with JetPrime reagents as per manufacturer’s instructions (Polyplus Transfection, NY). All transfections were for 48 hours unless otherwise noted.

### Reporter assays

The M-CSF promoter reporter (pMCSF-R-luc Addgene plasmid # 12420; http://n2t.net/addgene:12420; RRID: Addgene_12420)(Zhang et al., 1994) was used to assess RUNX1 transcriptional activity. 293T cells were transiently transfected with luciferase-based reporter plasmids and expression plasmids using jetPRIME (Polyplus, New York, NY). Total DNA quantity was constant across all wells. 48 hours post-transfection, luciferase assay reagent was mixed in a 1:1 ratio with cell lysate (Luciferase Assay System kit, Promega, Madison, WI). Luciferase activity was measured with Synergy H4 Hybrid Reader (BioTek, Winooski, VT). Transfection efficiency for each well was normalized using 62.5ng of a pCMV β-galactosidase plasmid, which was co-expressed in each experiment.

## Results

### hnRNP K is overexpressed in AML patient samples and correlates with poor clinical outcomes

We first assessed hnRNP K protein expression in AML using a publicly available reverse phase protein array (RPPA) dataset (Hu et al., 2019). While hnRNP K expression varied, AML cases had significantly higher median hnRNP K expression compared to healthy human bone marrow (p=0.0056, Fig 1A). As expected, a small percentage of cases had decreased hnRNP K expression, consistent with previous descriptions of *HNRNPK* haploinsufficiency corresponding with del(9q) (Gallardo et al., 2015; Kronke et al., 2013; Peniket et al., 2005). Increased hnRNP K expression was further associated with a statistically significant decrease in overall survival (OS; 24.3 months versus 48.7 months; HR 1.9; 95% CI 1.3-2.7; Figure 1B). Thus, stratification of patients based solely on hnRNP K overexpression was sufficient to elucidate a subset of patients with an inferior clinical outcome, suggesting that increased amounts of hnRNP K may be involved in the pathology of AML.

**Figure 1.**
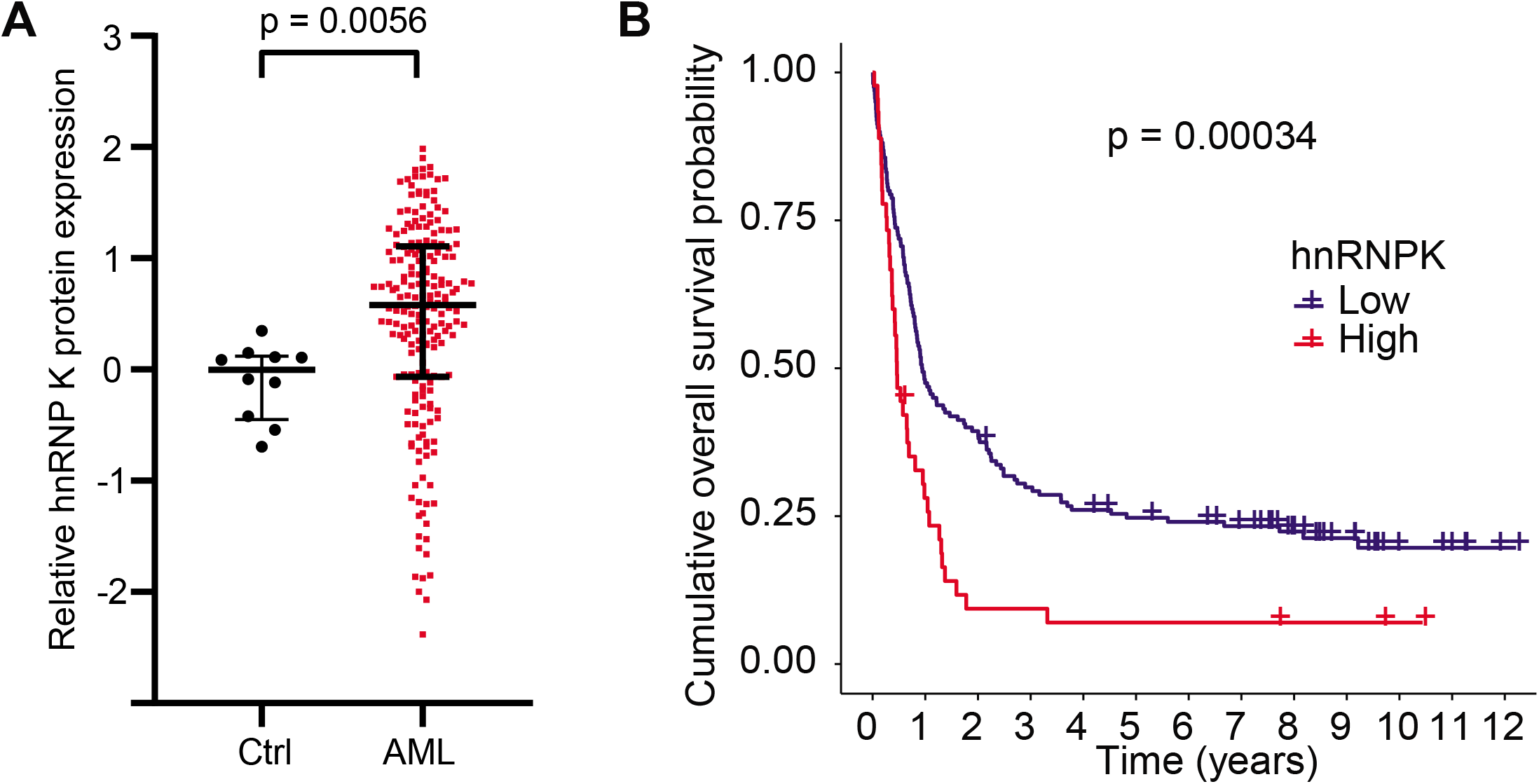
: hnRNP K expression in *de novo* AML. (A) Relative hnRNP K protein expression as quantified by RPPA. CD34+ cells from healthy human donor bone marrows (n=10) are indicated in black. Cells from bone marrows of patients with AML (n=205) are shown in red. (C) Overall survival of AML patients with either high hnRNP K protein expression as determined by RPPA (red; n=45) or normal/low hnRNP K expression (blue; n=160). Data is publicly available at www.leukemiaatlas.org.

We considered the possibility that increased levels of hnRNP K could be a consequence of mutation. In line with the findings of others, a mere 2.9% of AML cases at our institution were found to have an *HNRNPK* mutation (Supplemental Figure 1) (Cancer Genome Atlas Research et al., 2013; Papaemmanuil et al., 2016; Tyner et al., 2018). Given the discordance between this small mutational occurrence and the relatively large proportion of AML cases with elevated hnRNP K protein levels, we therefore posit that increased hnRNP K is of the wildtype form.

### hnRNP K overexpression impacts the differentiation potential of HSPCs

To evaluate a possible role of hnRNP K overexpression in myeloid disease, we overexpressed wildtype hnRNP K in murine hematopoietic stem and progenitor cells (HSPCs). Robust overexpression of hnRNP K was observed at the protein level (Figure 2A). *In vitro*, hnRNP K-overexpressing cells exhibited a significant increase in colony formation (p=0.008, Figures 2B-C), suggesting that elevated hnRNP K may influence self-renewal capacity of HSPCs. Flow cytometry revealed that hnRNP K-overexpressing colonies were composed of fewer Gr1^+^CD11b^+^ mature myeloid cells compared to controls (Figures 2D-E). However, immature c-kit^+^Sca-1^+^ cells were more prominent in hnRNP K-overexpressing colonies compared to controls (Figures 2D-E). These data suggested that hnRNP K overexpression hinders HSPC differentiation into mature myeloid cells and may be involved in myeloid leukemogenesis.

**Figure 2.**
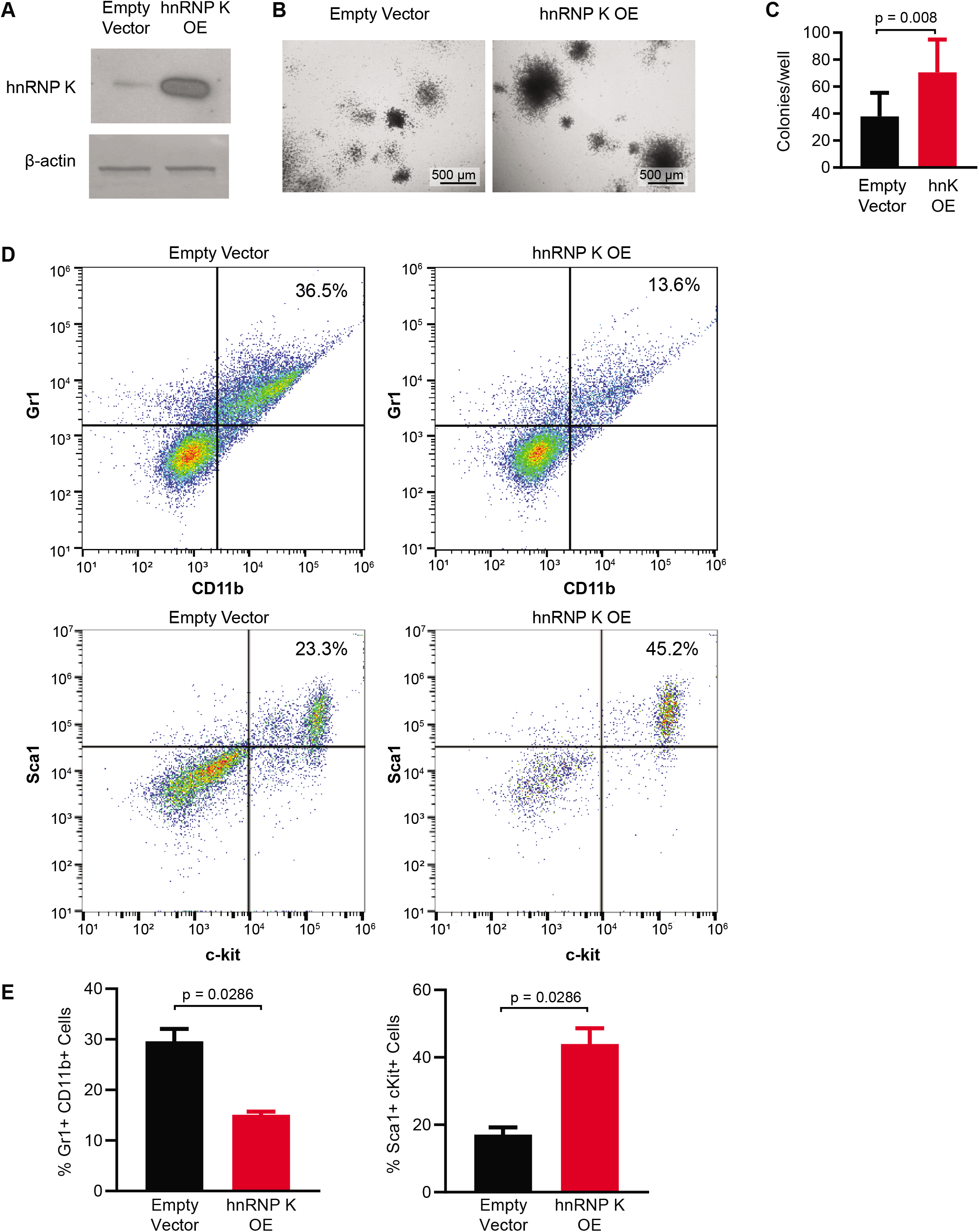
: hnRNP K overexpression in murine HSPCs. (A) Western blot showing hnRNP K protein levels in HSPCs infected with empty vector or hnRNP K plasmids. (B) Representative brightfield images of colonies from HSPCs infected with empty vector or hnRNP K plasmid. Scale bar represents 500 µm. (C) Bar graph quantitating the number of colonies per well in HSPCs infected with empty vector (n=14) and hnRNP K plasmids (n=10). (D) Flow cytometry analysis of HSPCs infected with empty vector and hnRNP K plasmids. (E) Bar graph quantitating the percentage of Gr1+CD11b+ and c-kit+Sca-1+ cells in HSPCs infected with empty vector (n=4) and hnRNP K plasmid (n=4).

### Overexpression of wildtype hnRNP K is sufficient to drive myeloid disease in mice

Because hnRNP K overexpression was observed in AML and correlated with poor clinical outcomes, we sought to directly evaluate if overexpression of wildtype hnRNP K was sufficient to drive myeloid disease. To this end, we injected murine HSPCs overexpressing wildtype hnRNP K or an empty vector control into sub-lethally irradiated NSG mice. Recipients of hnRNP K-overexpressing HSPCs had markedly shortened survival compared to recipients of control HSPCs (median survival 8.1 weeks versus median not reached (HR 3.0, 95% CI 1.2 – 7.3, p=0.02; Figure 3A), despite similar engraftment between groups (Figure 3B).

**Figure 3.**
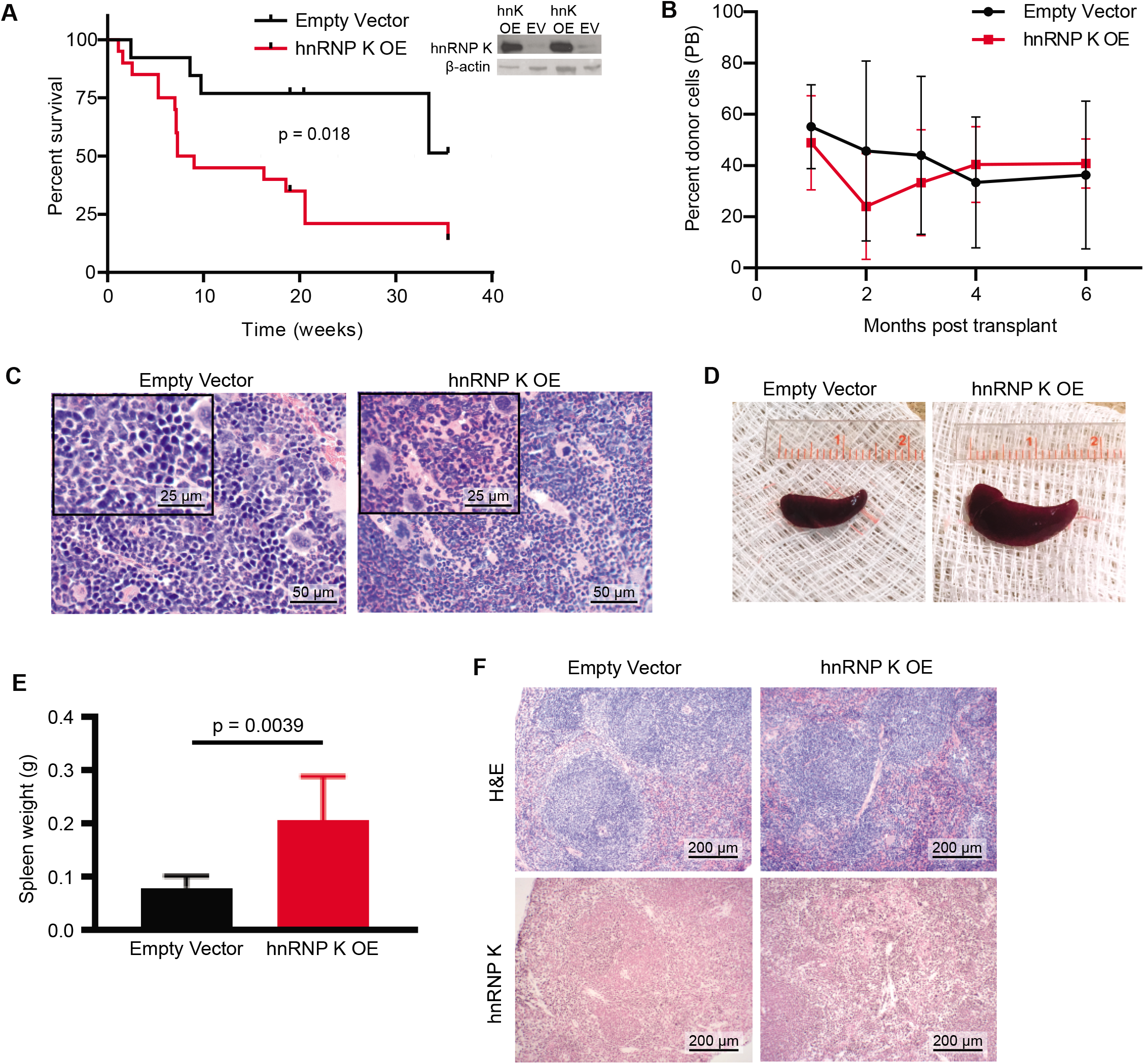
: Phenotypes observed in mice transplanted with hnRNP K overexpressing HSPCs. (A) Kaplan-Meier curves indicating survival of mice transplanted with HSPCs infected with an empty vector (n=13) and hnRNP K overexpression plasmid (n=22). Expression levels for hnRNP K in the HSPCs is depicted in the inset. (B) Percent donor cells in the peripheral blood of mice transplanted with HSPCs. (C) H&E staining in bone marrow samples obtained from mice transplanted with empty vector and hnRNP K overexpressing HSPCs. The scale bar represents 50 µm. (D) Representative photos of spleens from mice transplanted with empty vector and hnRNP K overexpressing HSPCs. (E) Bar graph depicting spleen weights from mice transplanted with empty vector (n=12) and hnRNP K overexpressing (n=21) HSPCs. (F) H&E and immunohistochemical analyses for hnRNP K in spleen samples from mice transplanted with empty vector or hnRNP K overexpressing HSPCs. The scale bar represents 200 µm.

Analysis of peripheral blood revealed that recipients of hnRNP K-overexpressing HSPCs had modest leukocytosis (Supplemental Figure 2A), while hemoglobin and platelet counts were not significantly changed between groups (Supplemental Figures 2B-C). Therefore, hnRNP K-overexpression appears to selectively affect the leukocyte compartment in our murine model. Indeed, mature neutrophils were less abundant in peripheral blood of mice transplanted with hnRNP K-overexpressing HSPCs (Supplemental Figure 2D). While the percentage of circulating monocytes or eosinophils did not differ between groups (Supplemental Figures 2E-F), hnRNP K-overexpression caused a slight increase in lymphocytes (Supplemental Figure 2G).

Given the peripheral blood abnormalities, we next evaluated bone marrow. Recipients of hnRNP K-overexpressing HSPCs had more cellular bone marrow than control recipients (Figure 3C). In addition, myeloid cells were overrepresented in bone marrow containing hnRNP K-overexpressing cells, and mild eosinophilia was observed (Figure 3C).

Given these abnormalities, we examined other hematopoietic organs. Splenomegaly was observed in recipients of hnRNP K-overexpressing HSPCs (Figure 3D-E) and histologic evaluation revealed starkly disrupted splenic architecture in recipients of hnRNP K-overexpressing HSPCs that corresponded with increased hnRNP K expression (Figure 3F). Immunohistochemistry on spleens from recipients of hnRNP K-overexpressing HSPCs showed CD34 and CD117 positive cells, indicating the presence of immature hematopoietic cells in this organ (Supplemental Figure 3A), which is in line with our observations from the colony formation assays.

Recipients of hnRNP K-overexpressing HSPCs also harbored infiltrations of leukocytes into the hepatic parenchyma that were not present in mice transplanted with empty vector-containing HSPCs (Supplemental Figure 3B). These hepatic infiltrates observed in recipients of hnRNP K-overexpressing HSPCs were largely negative for CD3, but positive for CD117, CD14, and myeloperoxidase (MPO; Supplemental Figure 3C), indicating that immature hematopoietic cells and cells of myeloid origin were present in this organ. Notably, lack of CD3 expression largely ruled out a graft versus host effect. Taken together, these data indicate that hnRNP K overexpression in hematopoietic stem and progenitor cells is sufficient to drive fatal myeloid disease in mice.

### hnRNP K binds to and influences the alternative splicing of the *RUNX1* transcript

Next, we wanted to understand a mechanistic basis for hnRNP K’s influence on myeloid development. To this end, we performed RNA-Seq on hnRNP K-overexpressing HSPCs. In addition to differences in gene expression (Supplemental Figure 4A), we observed statistically significant changes in numerous RNA splicing events (Figure 4A). Given the RNA-binding properties of hnRNP K coupled with the increasing appreciation for altered splicing events in hematologic malignancies, we focused on differentially spliced genes. When we specifically evaluated hematopoietic genes that were differentially spliced in the context of hnRNP K overexpression, we identified *Runx1* as one of the most significantly differentially spliced genes (Figure 4B). This was a particularly intriguing finding, as *RUNX1* was also among the transcripts most significantly associated with hnRNP K in an hnRNP K RNA-immunoprecipitation dataset in a human AML cell line generated from our lab (GSE 126479; Figure 4C (Gallardo et al., 2019)). These data, as well as the well-described role of RUNX1 as a critical mediator in many leukemias, led us to focus on studying this target.

**Figure 4.**
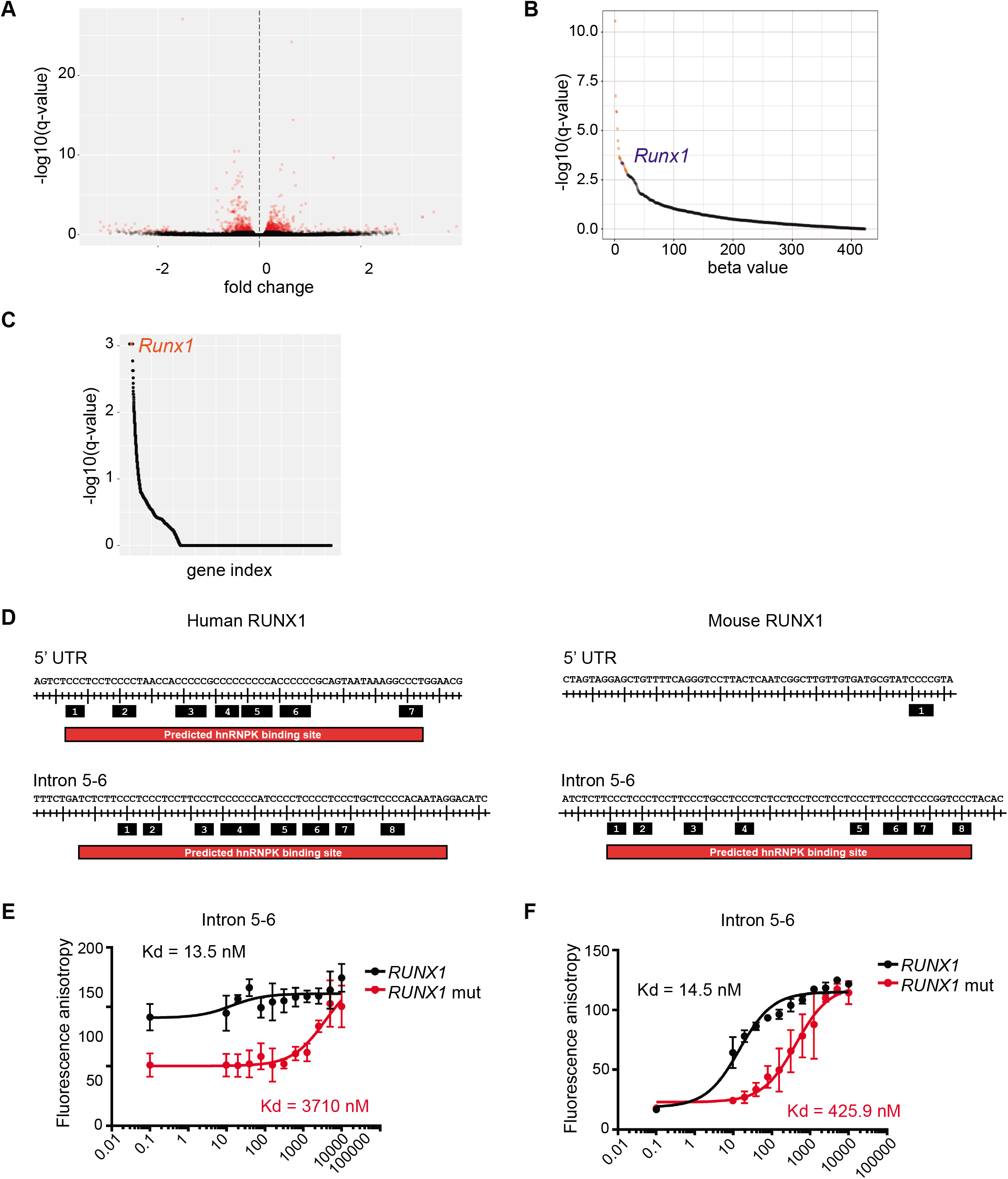
: Mechanistic basis for the oncogenicity of hnRNP K. (A) Volcano plot indicating alternatively spliced transcripts in HSPCs infected with empty vector (n=3) and hnRNP K plasmid (n=3). (B) Graph depicting differentially spliced transcripts in HSPCs infected with hnRNP K plasmid compared to empty vector, subsetted by causal implication in heme malignancies. (C) Graph depicting hnRNP K-associated transcripts, as determined by fRIP analysis, subsetted by causal implication in heme malignancies. The fRIP-Seq experiments were previously performed in the laboratory in triplicate in OCI-AML3 cells. (D) Identification of putative hnRNP K binding sites in the human and murine *RUNX1* transcripts using a previously generated computer algorithm. (E & F) Fluorescence anisotropy binding curves for purified full-length hnRNP K with FAM-labeled human and murine RUNX intron 5-6 wild-type and mutant oligos. Binding assays were performed in triplicate.

While hnRNP K has been associated with *Runx1* splicing (Cao et al., 2012), this has not been described in the context of hnRNP K overexpression, nor has a direct interaction between hnRNP K and *Runx1* RNA been characterized. Therefore, we used our previously developed computer algorithm to identify putative hnRNP K binding sites within human *RUNX1* or mouse *Runx1* transcripts (Gallardo et al., 2019). Two putative hnRNP K binding sites were identified in human *RUNX1*— one near the 3’ splice site of the intron 5-6 junction and one in the 5’ untranslated region (UTR) of *RUNX1b/c* (Figure 4D). Notably, only the intron 5-6 site was conserved in murine *Runx1*, which is consistent with a previous report of a similar site in the rat homolog (Cao et al., 2012).

To assess whether hnRNP K directly bound these sites in *RUNX1* RNA, we performed fluorescence anisotropy assays. Indeed, purified hnRNP K protein stringently bound the intron 5-6 site in both human and mouse *RUNX1* (Figures 4E-F). hnRNP K also bound tightly to the predicted site in the 5’ UTR of human *RUNX1* RNA (Supplemental Figure 4B). In all cases, the hnRNP K-*RUNX1* interaction was abrogated when the hnRNP K consensus binding site was mutated (Figures 4E-F, Supplementary Figure 4B), indicating that hnRNP K binds *RUNX1* directly and in a sequence-specific manner. These findings were confirmed in thermal shift assays, where hnRNP K was stabilized (i.e. had an increased melting temperature) in the presence of the conserved nucleic acid sequence corresponding to intron 5-6 in both human and murine *RUNX1* (Supplemental Figure 4C). As with the fluorescence anisotropy assays, mutations in the hnRNP K binding site diminished the ability of hnRNP K to be stabilized by these oligonucleotides (Supplemental Figure 4C).

Given the similarity between the hnRNP K binding site in *RUNX1* intron 5-6 in mouse and human, as well as RNA-Seq data identifying (Figure 4) that the *Runx1* splicing alterations surrounded exon 6, we focused our efforts on this site. Indeed, PCR amplification spanning exons 5 through 7 of *Runx1* in HSPCs validated the observations from RNA-Seq that an isoform of *Runx1* lacking exon 6 was enriched in hnRNP K-overexpressing HSPCs (Figure 5A). Sanger sequencing of a similar *RUNX1* PCR amplicon of exons 5-7 in human 293T cells further confirmed that the two PCR products amplified from this PCR contained exons 5-7 (referred to as FL *RUNX1;* 352 base pairs) or completely lacked exon 6 (referred to as *RUNX1ΔEx6;* 192 base pairs; Figure 5B). These results indicate that both *RUNX1 FL* and *RUNX1ΔEx6* isoforms are present in both human and mouse.

**Figure 5.**
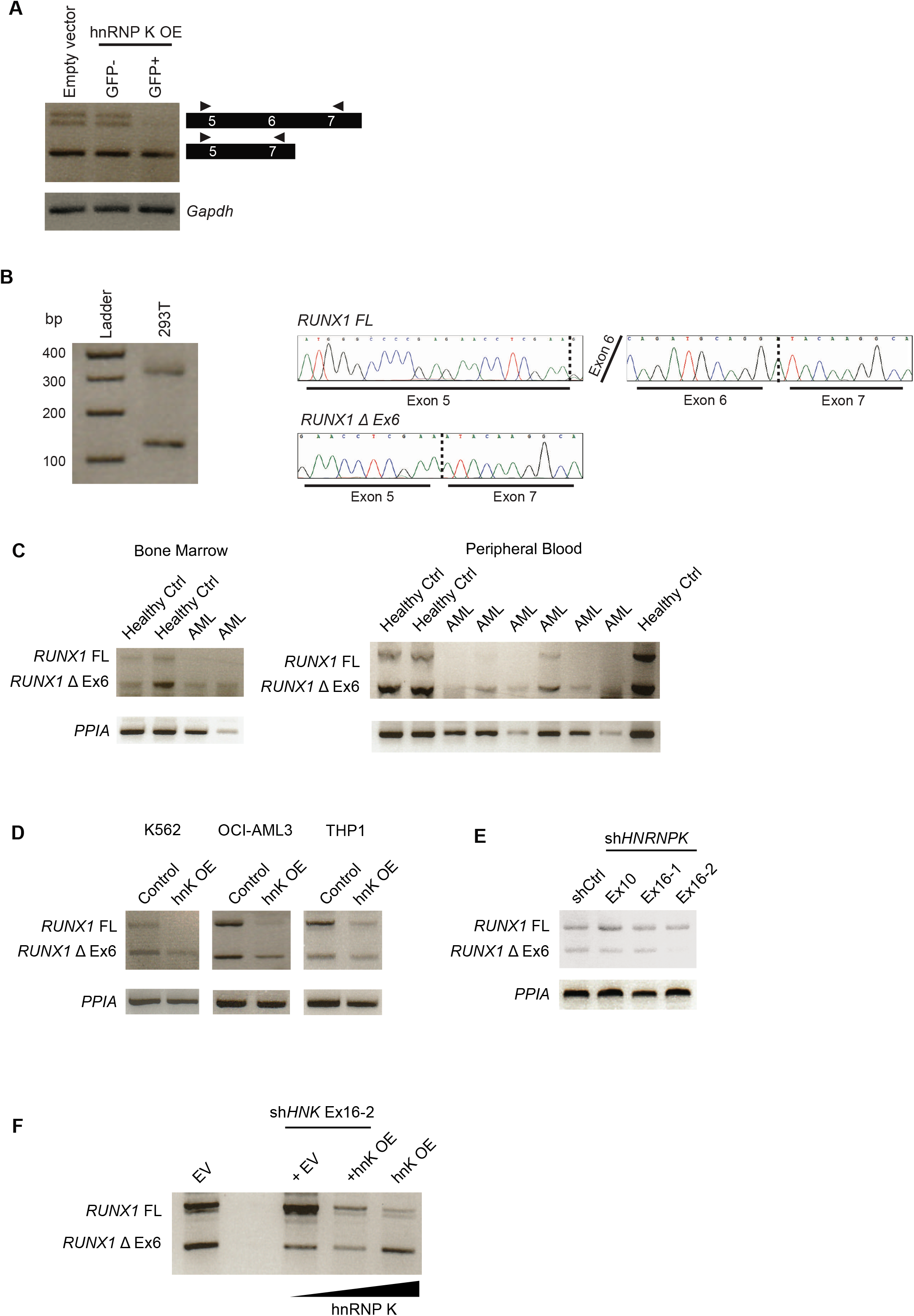
: Assessing hnRNP K’s impact on *RUNX1* alternative splicing. (A) RT-PCR based splicing analysis for *RUNX1* full-length and ΔEx6 isoforms in HSPCs infected with empty vector or hnRNP K plasmid. *Gapdh* was used as a loading control. (B) RT-PCR splicing analysis for *RUNX1* full-length and ΔEx6 isoforms in 293T cells followed by Sanger sequencing to confirm the sequences of the splice isoforms under study. (C) RT-PCR based splicing analysis for *RUNX1* full-length and ΔEx6 isoforms in peripheral blood and bone marrow cells obtained from healthy controls or AML patients. *PPIA* was used as a loading control. (D) RT-PCR based splicing analysis for *RUNX1* full-length and ΔEx6 isoforms in stable cell lines (K562, OCI-AML3, THP1) that overexpress hnRNP K. *PPIA* was used as a loading control. (E) RT-PCR based splicing analysis for *RUNX1* full-length and ΔEx6 isoforms in K562 stable cell lines with shRNA-mediated hnRNP K knockdown. One shRNA targets the hnRNP K CDS (Ex10) and two hairpins target the 3’-UTR (Ex16-1 and Ex16-2). PPIA was used as a loading control. (F) RT-PCR based splicing analysis for *RUNX1* full-length and ΔEx6 isoforms in K562 stable cell lines with shRNA-mediated hnRNP K knockdown rescued with hnRNP K overexpression. *PPIA* was used as a loading control.

### The *RUNX1ΔEx6* splice isoform is present in AML patient samples

While these isoforms of *RUNX1* were present in human and mouse, we wanted to examine whether these were also present in human AML samples. Indeed, *RUNX1ΔEx6* was prominent in bone marrow and peripheral blood of patients with AML (Figure 5C). Analysis of publicly available data also corroborated the presence of this *RUNX1ΔEx6* in AML (Supplementary Figure 5A). This suggests that alterations in *RUNX1* splicing involving exon 6 inclusion are clinically evident and may be relevant.

### hnRNP K expression levels regulate the generation of the *RUNX1ΔEx6* splice isoform

To explicitly understand how hnRNP K levels mediate this splicing event, we developed several inducible hnRNP K-overexpressing human leukemia cell lines (Supplemental Figure 5B). In each case, cells that overexpressed hnRNP K had a relative enrichment of *RUNX1ΔEx6* at the expense of *RUNX1 FL* compared to cells expressing an empty vector control (Figure 5D). Strikingly, knockdown of hnRNP K induced the opposite effect, as cells transduced with sh*HNRNPK* had increased *RUNX1 FL* at the expense of *RUNX1ΔEx6* (Figure 5E and Supplementary Figure 5C). Importantly, re-introduction of hnRNP K into the sh*HNRNPK-*knockdown cells rescued the relative proportion of *RUNX1 FL* to *RUNX1ΔEx6* to near that of controls (Figure 5F and Supplementary Figure 5D). This indicates that inclusion or exclusion of *RUNX1* exon 6 is dependent on hnRNP K levels. Importantly, this observation extended to the protein expression of RUNX1 as well, where the shorter isoform of RUNX1 was detectable at higher levels in the presence of hnRNP K overexpression (Supplemental Figure 5E). In contrast, knockdown of hnRNP K resulted in a reduction in the expression of this shorter RUNX1 isoform (Supplemental Figure 5F).

### The *RUNX1ΔEx6* splice isoform has differential protein stability and function

Since hnRNP K overexpression leads to relative enrichment of *RUNX1ΔEx6*, we next sought to evaluate any functional consequences of lacking this exon. Cycloheximide chase assays, in cells with stable inducible expression of RUNX1 FL or RUNX1ΔEx6, demonstrated that RUNX1ΔEx6 was substantially more stable than RUNX1 FL, in line with previous observations (Figure 6A) (Komeno et al., 2014; Sun et al., 2020). This effect was almost completely abrogated by addition of MG132 (Figure 6B), indicating that exon 6 is a critical mediator of RUNX1 protein stability. This is consistent with reports that have identified ubiquitination sites in exon 6 of the RUNX1 protein (Biggs et al., 2006).

**Figure 6.**
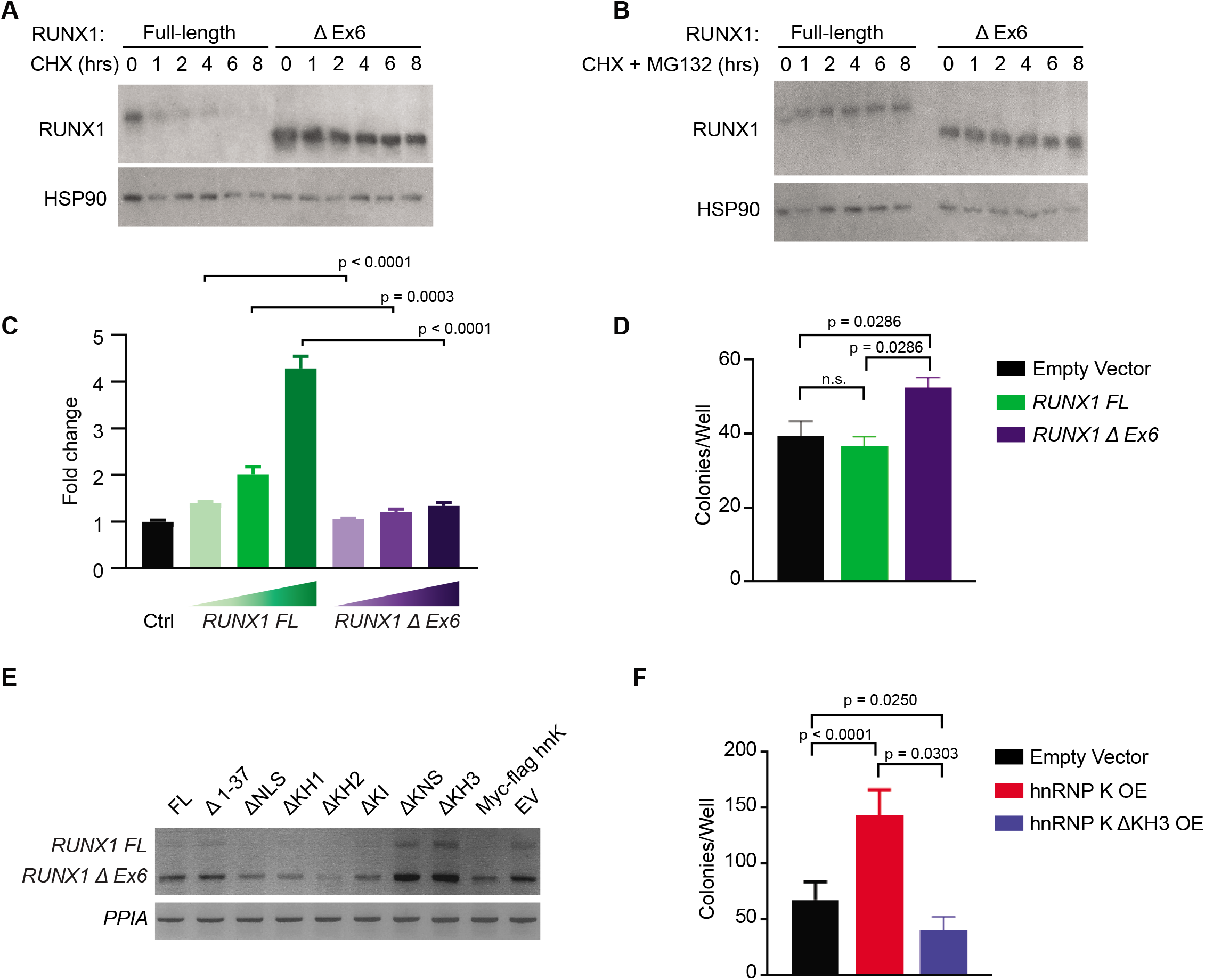
: Functional relevance of the *RUNX ΔEx6* splice isoform. (A) Immunoblot for RUNX1 following a cycloheximide chase experiment in 293T cells that stably express *RUNX1* full-length or RUNX ΔEx6. HSP90 serves as the loading control. (B) Immunoblot for RUNX1 following cycloheximide+MG132 treatment in 293T cells that stably express *RUNX1* full-length or RUNX ΔEx6. HSP90 serves as the loading control. (C) Luciferase-based reporter assay to assess transactivation from a CSF-1R promoter reporter in 293T cells. The experiment was independently done three times in triplicate. (D) Bar graph representing number of colonies formed per well for HSPCs infected with empty vector (n=7), *RUNX1* full-length (n=4) or RUNX ΔEx6 (n=4) plasmid. (E) RT-PCR based splicing assay (in 293T cells) to determine the impact of different hnRNP K protein domains in regulating RUNX1 alternative splicing. (F) Bar graph representing number of colonies formed per well for HSPCs infected with empty vector (n=7), hnRNP K (n=7) or hnRNP K ΔKH3 (n=3) plasmid.

As RUNX1 is a well-defined transcription factor, we then queried whether lack of exon 6 would affect the transcriptional capabilities of this protein. Indeed, luciferase assays indicated that RUNX1 FL and RUNX1ΔEx6 had differential abilities to transactivate a promoter-reporter derived from the CSF1R promoter, which is known to have RUNX1 binding sites (Figure 6C). Together, our data demonstrate that RUNX1 FL and RUNX1ΔEx6 are phenotypically and functionally distinct isoforms.

To understand whether this isoform of RUNX1 may be responsible, in part, for mediating the hnRNP K-overexpression phenotype, we repeated colony formation assays using HSPCs. HSPCs overexpressing RUNX1ΔEx6 formed significantly more colonies *in vitro* than HSPCs overexpressing RUNX1 FL or an empty vector (Figure 6D). This increase in colony formation was similar to that observed with overexpression of hnRNP K (Figure 2B-C), suggesting that increased expression of RUNX1ΔEx6 may be partly responsible for this hnRNP K-mediated effect.

### The KH3 RNA-binding domain of hnRNP K is required for the generation of the *RUNX1ΔEx6* splice isoform

Given the striking splicing differences observed with hnRNP K overexpression in both human and mouse, we next set out to determine which domain of the hnRNP K protein was responsible for mediating this splicing event as this could expedite future drug discovery efforts. When hnRNP K was overexpressed, the relative proportion of *RUNX1 FL* to *RUNX1ΔEx6* was decreased, as we had observed in earlier studies (Figure 6E and Supplemental Figure 6B). However, when hnRNP K lacking either the KNS or KH3 domain was overexpressed, this effect was largely mitigated, and the *RUNX1* splicing pattern resembled control cells (Figure 6E and Supplemental Figure 6). This suggests that isolated domains of hnRNP K (KH3 and possibly KNS) are responsible for the splicing surrounding RUNX1 exon 6. Given that the splicing process occurs in the nucleus and requires direct RNA-binding activity, it is therefore understandable why either the RNA-binding KH3 domain or the nuclear localization KNS domain would be critical.

Since the KH3 domain of hnRNP K appeared to be largely responsible for the expression of *RUNX1ΔEx6*, we overexpressed the isoform of hnRNP K lacking KH3 in HSPCs and performed colony formation assays. Strikingly, the absence of KH3 completely reversed the increase in colony formation associated with overexpression of full-length hnRNP K (Figure 6F). This demonstrates that the KH3 domain is critical not only for splicing surrounding exon 6 of *RUNX1*, but also that the KH3 domain is required for hnRNP K to exert its pro-growth activity in HSPCs. Taken together, these data demonstrate that the RNA-binding protein hnRNP K is highly expressed in AML. Overexpression of wildtype, unmutated hnRNP K increases colony formation potential of HSPCs, while inhibiting terminal differentiation. These findings are echoed *in vivo*, where increased hnRNP K expression leads to a fatal myeloid disease. Mechanistically, hnRNP K directly binds *RUNX1* RNA in mouse and in human, and its overexpression results in an enrichment of the phenotypically and functionally distinct isoform of RUNX1 lacking exon 6 (*RUNX1ΔEx6*). Going forward, development of drugs to disrupt hnRNP K’s interaction with RNA, perhaps by specifically targeting the KH3 domain of hnRNP K, may have therapeutic potential in hnRNP K-overexpressing malignancies, including AML.

## Discussion

RNA-binding proteins (RBPs) are increasingly being considered as essential molecules in regulating normal and malignant hematopoiesis. Spatio-temporal regulation of the gene targets of RBPs allows for the development and function of different hematological lineages (Kharas et al., 2010a; Wang et al., 2019). Consequently, aberrant function of RBPs, either due to mutations or altered expression, results in pathogenic states in the hematopoietic system (Barbieri et al., 2017; Gallardo et al., 2019; Kharas et al., 2010a; Vu et al., 2017). In this work, we describe overexpression of the RNA-binding protein, hnRNP K, in AML.

hnRNP K is frequently altered in malignancies and other disease states. Mutations in hnRNP K are relatively rare in hematological malignancies, as assessed by our patient cohort at MD Anderson Cancer Center (Supplemental Figure 1) and published work by other groups (Cancer Genome Atlas Research et al., 2013; Papaemmanuil et al., 2016; Tyner et al., 2018). Deletions of chromosome 9q, which encompass the *HNRNPK* locus at 9q21.32, are also rare events in AML (Gallardo et al., 2015; Kronke et al., 2013; Sweetser et al., 2005), but there has been sufficient evidence to establish the role of hnRNP K as a contextual haploinsufficient tumor suppressor. On the other hand, clinical and biological studies from our group and others have also identified that that *increased* hnRNP K expression may play a role in both hematologic malignancies and solid tumors (Barboro et al., 2009; Carpenter et al., 2006; Chen et al., 2010; Wen et al., 2010; Zhou et al., 2010). Indeed, increased expression of hnRNP K appears to be the more common occurrence in blood cancers (Figure 1A, (Gallardo et al., 2019)).

In the present work, using a previously published database, we observe that hnRNP K is overexpressed in patients with AML and correlates with poor clinical outcomes. Given the low prevalence of hnRNP K mutations in AML, we posit that it is the wildtype form of hnRNP K that is overexpressed. Further, molecular mechanisms that underlie the overexpression of hnRNP K in AML remain an active area of investigation in our laboratory.

To evaluate the direct role of hnRNP K in driving myeloid phenotypes, we used a murine HSPC transplant model. Transplantation of hnRNP K-overexpressing HSPCs into mice results in a fatal myeloproliferative phenotype. The aberrant myelopoiesis observed in our mouse model is intriguing for several reasons. We recently described a mouse model wherein B-cell lymphomas develop secondary to hnRNP K overexpression in B-cells (Gallardo et al., 2019). In the current study, where hnRNP K was overexpressed in HSPCs, no lymphomas were observed. This suggests that hnRNP K overexpression earlier in hematopoiesis promotes a myeloid bias and/or inhibits lymphoid differentiation. Consistent with this, microarray data in human hematopoiesis has revealed *HNRNPK* transcript expression is higher in myeloid-biased compared to lymphoid-biased progenitor cells (Novershtern et al., 2011; Verhaak et al., 2009). In addition, we observed similar phenotypes including shortened survival and myeloid hyperplasia in mice that are haploinsufficient for *Hnrnpk (Gallardo et al*., *2019)*, suggesting that normal hematopoiesis, particularly of the myeloid lineage, requires exquisitely tight regulation of hnRNP K expression.

Mechanistic studies reveal that hnRNP K overexpression results in altered pre-mRNA splicing events. One of the top genes that was differentially spliced was *RUNX1*, and we decided to focus on altered splicing of the *RUNX1* transcript given its critical role in myeloid biology. Specifically, overexpression of hnRNP K resulted in the preferential splicing and generation of a *RUNX1* isoform lacking exon 6. Even a small increase in the levels of the *RUNX1ΔEx6* isoform is likely to be physiologically relevant, given the drastically increased stability of the corresponding protein. The *Runx1ΔEx6* isoform exhibits higher self-renewal capacity in colony formation assays, which is in agreement with the findings of Komeno et al and Sun et al (Komeno et al., 2014; Sun et al., 2020). In addition, others have demonstrated a depletion in the cellular pool of HSCs *in vivo* in mice lacking the *RUNX1ΔEx6* isoform, indicating the importance of tight regulation of splicing in hematopoiesis (Ghanem et al., 2018; Komeno et al., 2014).

Functionally, we observed that the RUNX1ΔEx6 isoform has different transcriptional activities compared to the full-length isoform. This may be attributed to the fact that the protein domain encoded by exon 6 is responsible for the interaction between RUNX1 and the transcriptional co-repressor SIN3A (Zhao et al., 2008). The region encoded by exon 6 also contains arginine methylation sites, and these post-translational modifications have been implicated in regulation of transcriptional activity (Zhao et al., 2008). The exact mechanisms by which the RUNX1ΔEx6 isoform is generated and the molecular processes that impart functional diversity to this alternatively spliced isoform remains an active area of investigation. However, given the robust presence of this isoform in AML patient samples, it is likely to have a role in AML biology.

Taken together, our data demonstrate the pathogenicity of overexpression of a wild type RBP, hnRNP K, in AML. While the altered splicing of the *RUNX1* transcript is observed upon hnRNP K overexpression, it is unlikely to be the sole mechanism driving the oncogenicity of hnRNP K. However, this paper provides evidence that aberrant expression of an RBP can disrupt splicing within the cell and has far reaching cellular implications, including but not limited to altered transcription factor activities.

## Acknowledgements

We would like to thank Kendra Allton, Sabrina Stratton, and other members of Dr. Michelle Barton’s laboratory for helpful conversations, and Taghi Manshouri for technical assistance and collaborative efforts.

