## Supplemental Figure Legends for "Heterogeneous nuclear ribonucleoprotein K is overexpressed in acute myeloid leukemia and causes myeloproliferative disease in mice via altered *Runx1* splicing"

Supplemental Figure 1: Schematic indicating hnRNP K mutations occurring in AML patients at MD Anderson Cancer Center. Numbers indicate amino acid positions flanking functional domains, though diagram, though diagram is not to scale. Each mutation was observed one time.

Supplemental Figure 2: Analysis of peripheral blood of mice transplanted with HSPCs. Complete blood analysis was done for mice transplanted with HSPCs infected with either empty vector or an hnRNP K plasmid to determine: (A) WBC count, (B) hemoglobin, (C) platelet count, (D) segmented cell count, (E) monocyte count, (F) eosinophil count, and (G) lymphocyte count.

Supplemental Figure 3: Pathological analyses of spleen and liver samples from mice transplanted with HSPCs. (A) H&E staining and immunohistochemical analysis for CD34 and CD117 in spleens of mice transplanted with HSPCs. Scale bar represents 200 µm for low magnification and 50 µm for high magnification. (B) H&E staining of liver infiltrates from mice transplanted with HSPCs. The scale bar represents 200 µm. (C) H&E staining and immunohistochemical analysis for CD3, CD34, CD117 and MPO in liver infiltrates in mice transplanted with hnRNP K OE HSPCs. The scale bar represents 50 µm.

Supplemental Figure 4: Elucidating the molecular basis for the oncogenicity of hnRNP K. (A) Heat map depicting differentially expressed transcripts in HSPCs infected with empty vector or hnRNP K plasmids (n=3). (B) Fluorescence anisotropy binding curves for purified full-length hnRNP K with FAM-labeled human *RUNX1* 5’-UTR wild type and mutant oligos. Binding assays were performed in triplicate. (C) Bar graph depicting melting temperature of purified full-length hnRNP K protein with oligos derived from the human *RUNX1* transcript (wild type and mutant). Melting temperatures were determined using SYPRO orange dye in a thermofluor assay. At least 4 replicates were done for each experimental condition.

Supplemental Figure 5. Impact of hnRNP K on *RUNX1* splicing. (A) Levels of *RUNX1* full-length and ΔEx6 transcripts in AML patients from the BEAT AML 1.0 cohort (Tyner et al., 2018). Each dot represents an individual patient. Cases with reported *RUNX1* alterations are indicated in red (mutation) or blue (translocation). (B) Immunoblot analyses for hnRNP K in cell lines stably expressing hnRNP K (K562, OCI-AML3, THP1). β-actin is used as a loading control. (C) Immunoblot analyses for hnRNP K in K562 cell with stable knockdown of hnRNP K using three unique hairpins, Ex10, Ex16-1 and Ex 16-2. β-actin is used as a loading control. (D) Immunoblot analyses for hnRNP K in K562 stable cells with shRNA 16-2 mediated hnRNP K knockdown rescued with hnRNP K overexpression. β-actin was used as a loading control. (E) Immunoblot analyses for hnRNP K and RUNX1 in stable cell lines (K562 and OCI-AL3) overexpressing hnRNP K. β-actin was used as a loading control. (F) Immunoblot analyses for hnRNP K and RUNX1 in OCI-AML3 stable cell line with shRNA mediated hnRNP K knockdown.

Supplemental Figure 6. Immunoblot analysis for flag-tagged hnRNP K deletion plasmids transiently transfected in 293T cells. HSP90 was used as a loading control.
