## Supplementary figures and images for "Heterogeneous nuclear ribonucleoprotein K is overexpressed in acute myeloid leukemia and causes myeloproliferative disease in mice via altered *Runx1* splicing"

### Supplemental Figure 1

Aitken, M. et al. Supplemental Figure 1

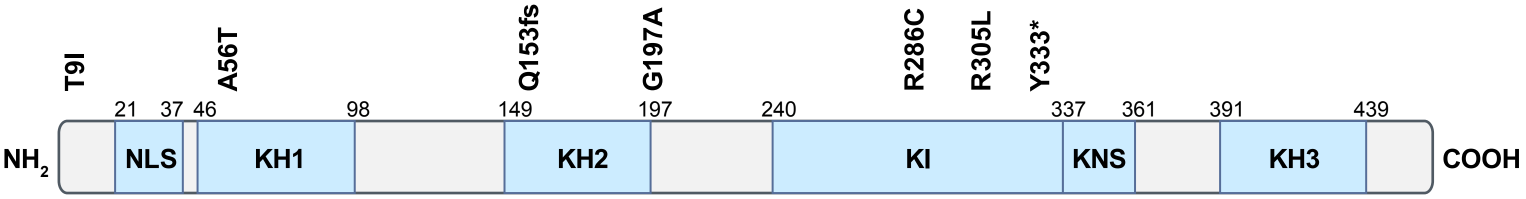

### Supplemental Figure 2

Aitken, M. et al. Supplemental Figure 2

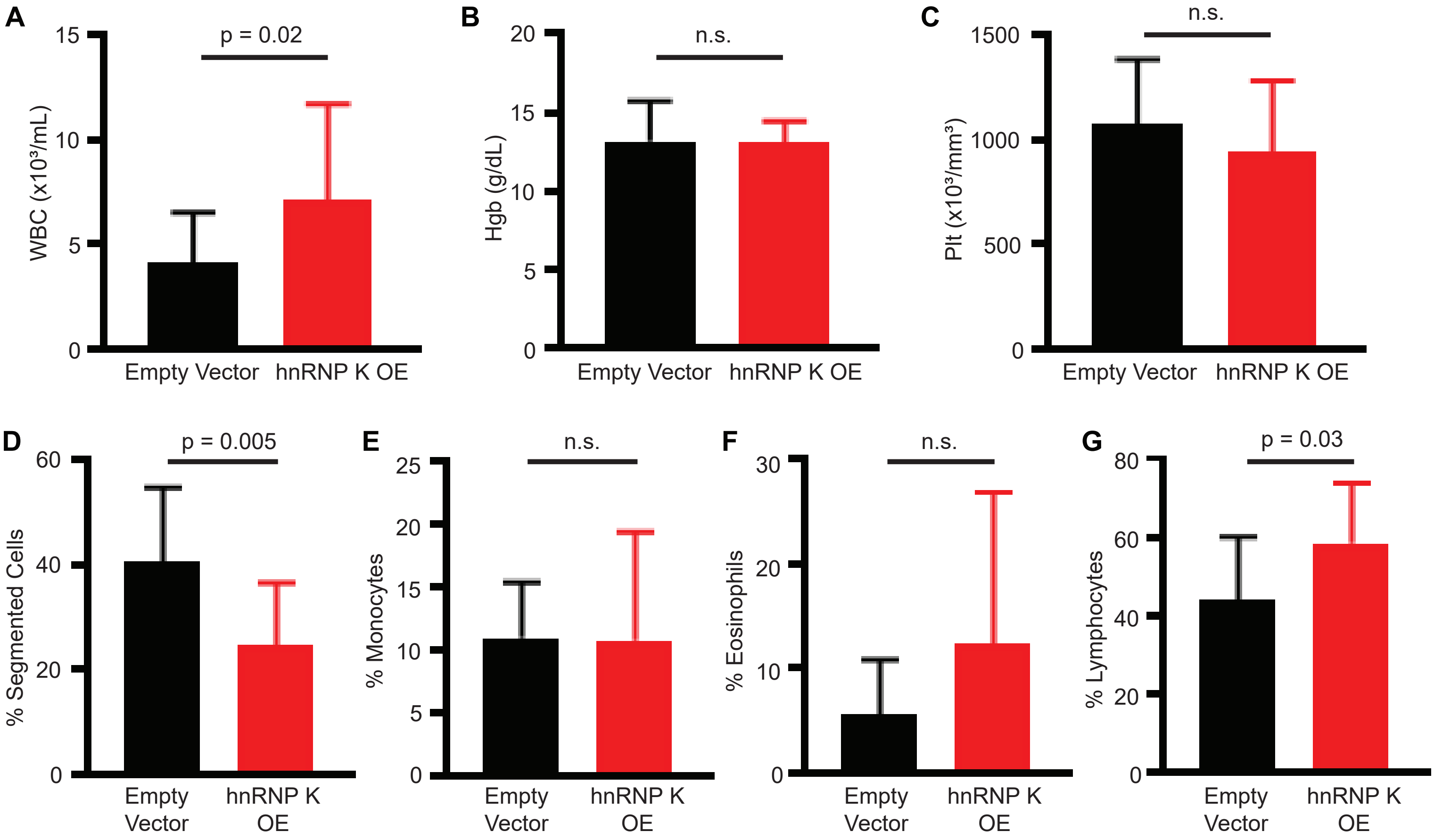

### supplemental Figure 3

Aitken, M. et al. Supplemental Figure 3

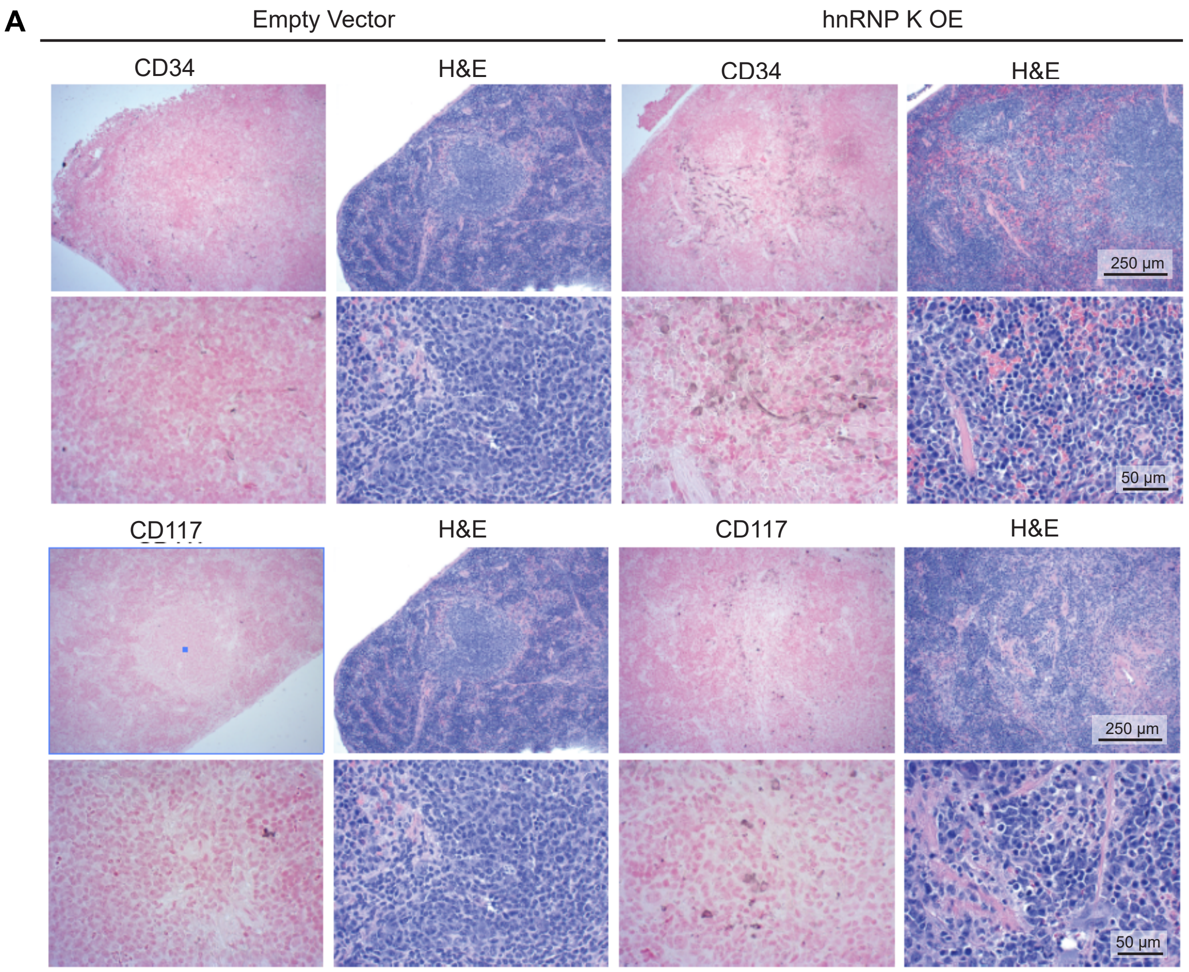


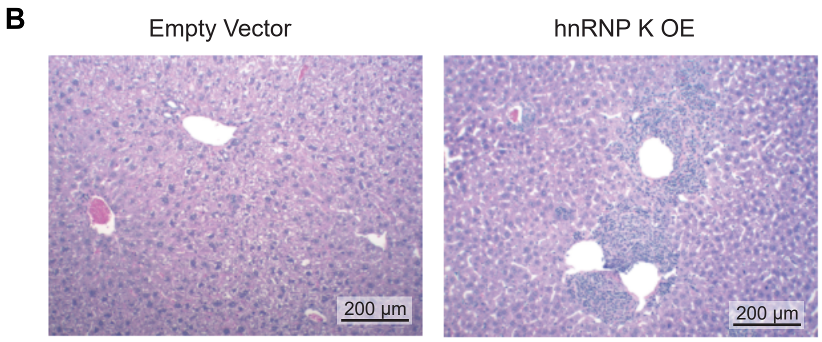


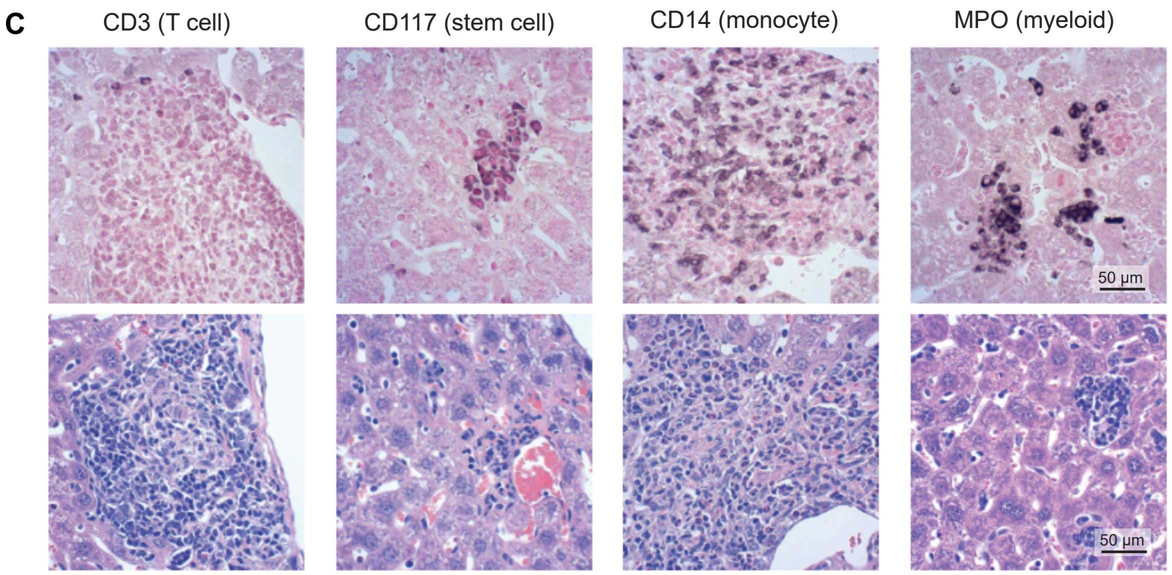

### Supplemental Figure 4

Aitken, M. et al. Supplemental Figure 4

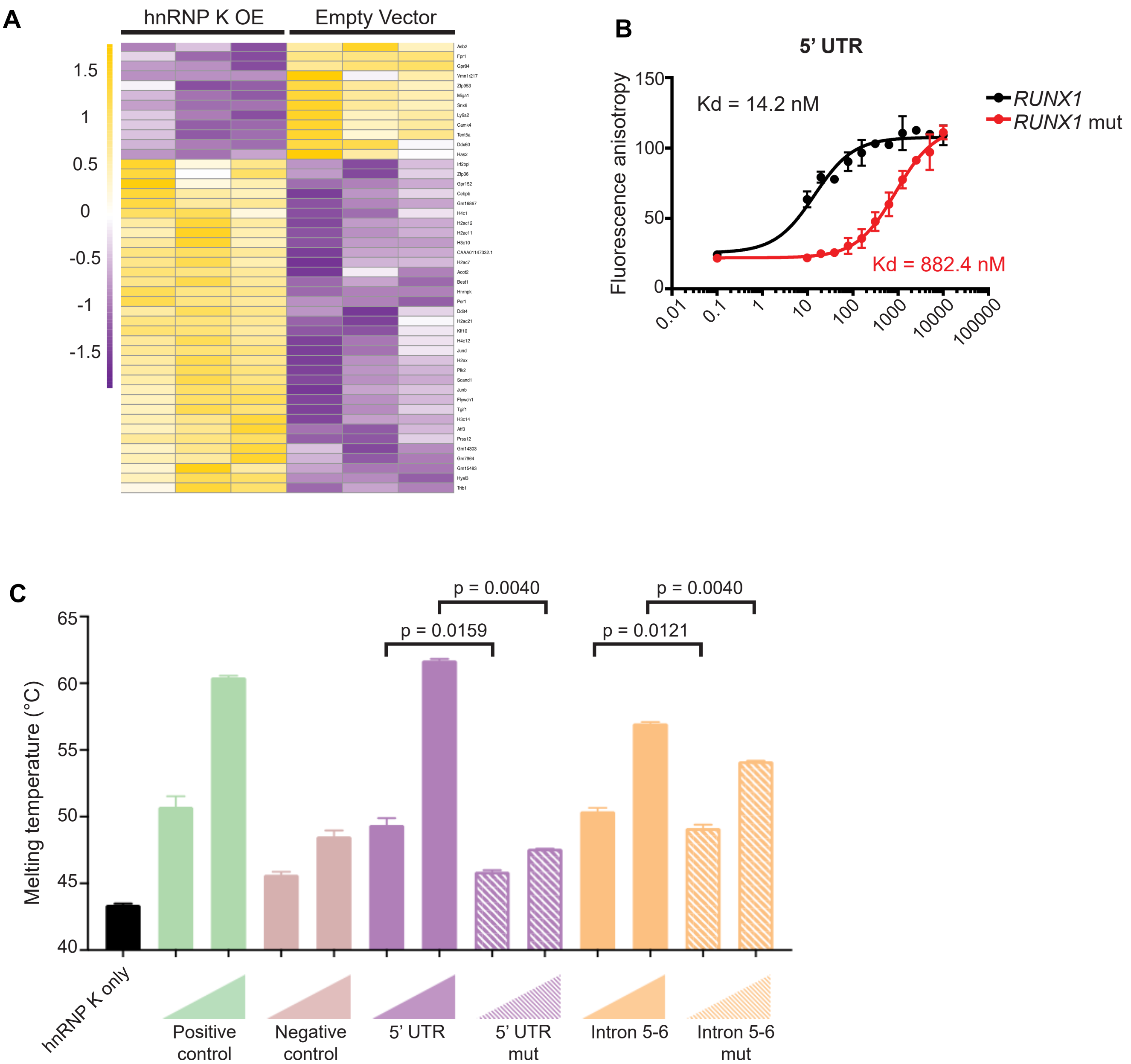

### Supplemental Figure 5

Aitken, M. et al. Supplemental Figure 5

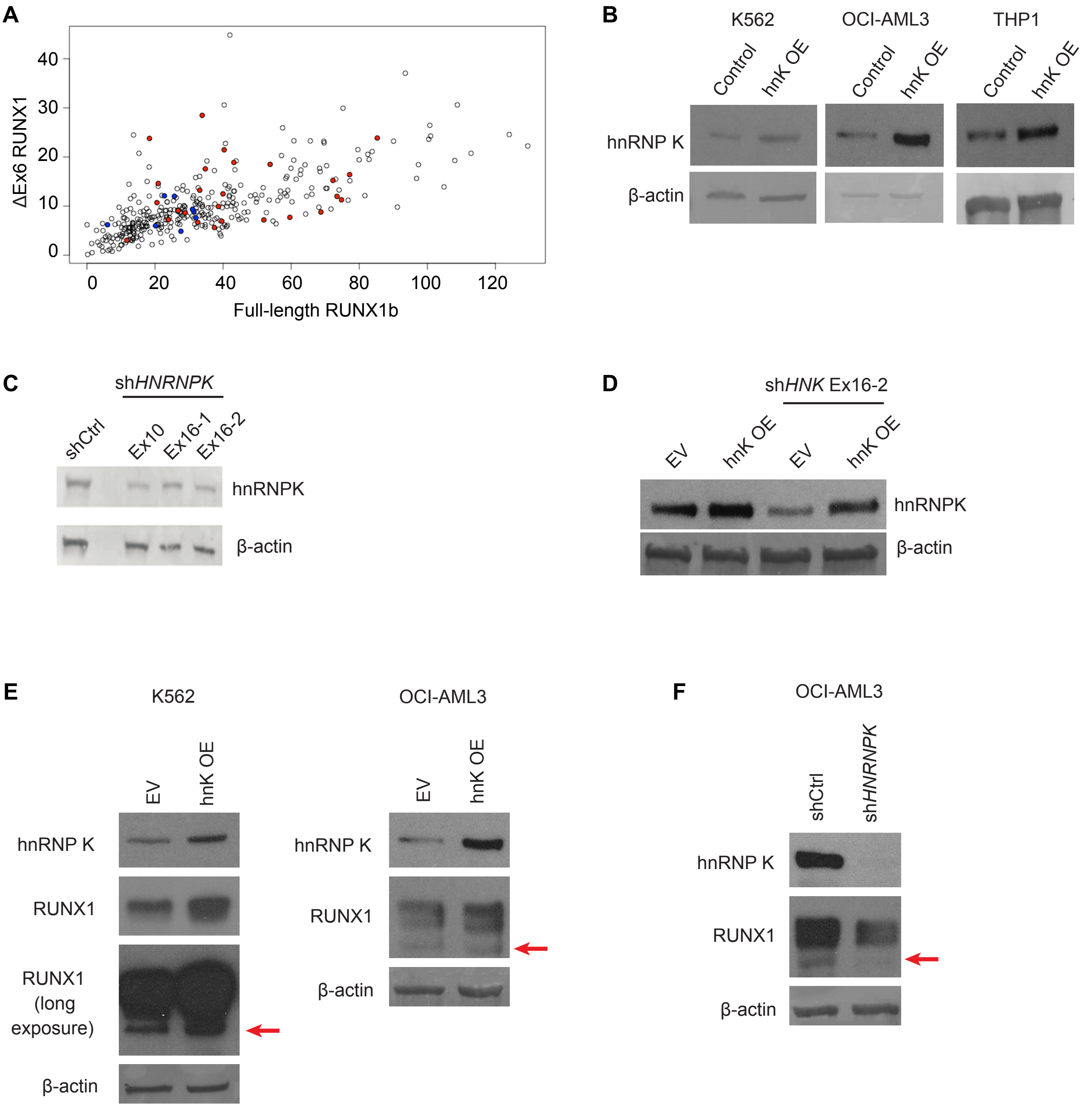

### Supplemental Figure 6

Aitken, M. et al. Supplemental Figure 6

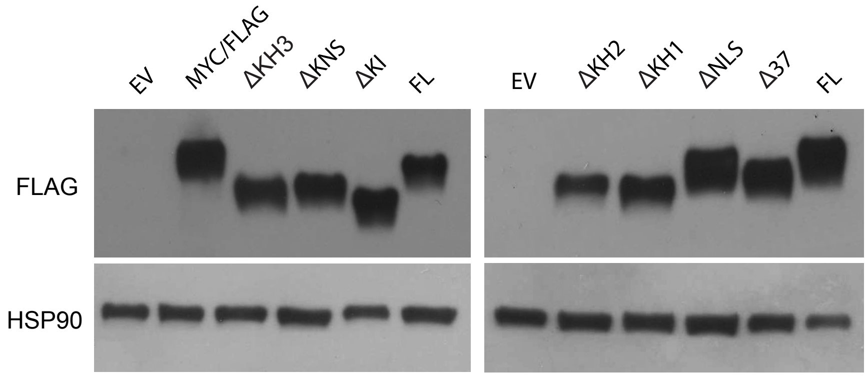
